# Histone modifications and DNA methylation act cooperatively in regulating symbiosis genes in the sea anemone Aiptasia

**DOI:** 10.1101/2022.05.10.491282

**Authors:** Kashif Nawaz, Maha J. Cziesielski, Kiruthiga G. Mariappan, Guoxin Cui, Manuel Aranda

## Abstract

The symbiotic relationship between cnidarians and dinoflagellates is one of the most widespread endosymbiosis in our oceans and provides the ecological basis of coral-reef ecosystems. Although many studies have been undertaken to unravel the molecular mechanisms underlying these symbioses, we still know little about the epigenetic mechanisms that control the transcriptional responses to symbiosis. Here, we used the model organism *Exaiptasia diaphana* to study the genome-wide patterns and putative functions of the histone modifications H3K27ac, H3K4me3, H3K9ac, H3K36me3 and H3K27me3 in symbiosis. While we find that their functions are generally conserved, we observed that colocalization of more than one modification and or DNA methylation correlated with significantly higher gene expression, suggesting a cooperative action of histone modifications and DNA methylation in promoting gene expression. Analysis of symbiosis genes revealed that activating histone modifications predominantly associated with symbiosis induced genes involved in glucose metabolism, nitrogen transport, amino acid biosynthesis and organism growth while symbiosis suppressed genes were involved in catabolic processes. Our results provide new insights into the mechanisms of prominent histone modifications and their interaction with DNA methylation in regulating symbiosis in cnidarians.

## Introduction

Coral reefs are often considered the rainforests of the sea, as they form marine-biodiversity hotspots. Reef ecosystem health directly depends on symbiotic cnidarians, such as corals and anemones, that provide essential habitats for a myriad of marine organisms. To thrive in the oligotrophic environment of tropical oceans, corals, and other symbiotic cnidarians, depend on an intimate endosymbiosis with photosynthetic dinoflagellates of the family *Symbiodiniaceae*, also known as zooxanthellae ^1–3^. Living within the host’s gastrodermal cells, the symbionts provide their hosts with over 90% of their total energy demands ^1^, making these symbiotic relationships vital for the functioning of the coral reef ecosystem. The disruption of this host-symbiont relationship, also known as bleaching, can result in extensive mortality and subsequent degradation and loss of entire coral reefs ^2–4^. Significant efforts have been made to understand the molecular mechanism underlying this relationship ^5^. However, there are still substantial knowledge gaps in our understanding of the molecular underpinnings of these relationships, especially pertaining to the role of epigenetic mechanisms in regulating the interactions between the host and the symbionts, which remain elusive in cnidarian symbiosis research ^6^. The uptake and maintenance of symbionts require specific host responses, such as the suppression of the immune system ^7, 8, 10, 15^ and the regulation of nutrient fluxes to control symbiont proliferation ^9^, to maintain a stable symbiotic relationship. Such responses are mediated through transcriptional changes that are known to be regulated via epigenetic mechanisms in other organisms ^10–14^, and some endosymbionts have even been shown to evoke such responses by directly modifying the epigenome of their hosts ^10, 15^. However, while many recent studies have highlighted the importance of epigenetic mechanisms in maintaining symbiotic relationships in plants and animals ^16–18^, only one study looking at the role of DNA methylation in symbiosis has been conducted in zooxanthellate cnidarians ^6^.

Eukaryotic genomes are packaged in the form of a DNA-protein complex termed chromatin. The structural unit of chromatin is known as the nucleosome, which consists of a core protein octamer and a stretch of ∼147 bp of DNA that is wound around it. The protein octamer comprises two of each of the core histones H2A, H2B, H3 and H4 and a non-core linker histone, histone H1, which provides external stability to nucleosomes ^19, 20^. This essential organization of histones aids in the folding of the DNA into a higher-order structure termed as chromatin fibers ^19–21^. The N-terminal tail of histone proteins can include reversible covalent changes termed post-translational modifications (PTMs), which control chromatin structure and, thus, the epigenetic regulation of gene expression and genome stability. These modifications have collectively been termed the histone code ^19^. Along with these modifications, other epigenetic factors such as DNA methylation and small RNAs collectively influence the chromatin structure, and in succession, the accessibility of the genetic information ^19, 20^.

Histone modifications can affect gene regulation differently depending on their type and location in the genome (Table 1). Among the different PTMs, acetylation and methylation of specific histone tail residues have been most extensively studied ^11–12^, and they have been found to promote repressing and activating roles in the regulation of gene expression ^13^. In general, activator complexes methylate or acetylate specific amino acid residues in tails of histones bound to gene promoter regions, thereby destabilizing the nucleosome-DNA interaction, and facilitating the assembly of the transcriptional machinery at the promoter. However, repressor complexes demethylate/deacetylate histone tails and strengthen the DNA histone interaction, resulting in hindered accessibility of the respective genomic regions for the transcriptional machinery ^14^.

**Table 1:** Histone modifications with their function and location.

| Histone modifications | Function | Location |
| --- | --- | --- |
| H3K4me3 | Activation | Promoters, bivalent domains |
| H3K36me3 | Activation | Gene-body |
| H3K9ac | Activation | Enhancers, promoters |
| H3K27ac | Activation | Enhancers, promoters |
| H3K27me3 | Repression | Promoters in gene-rich regions, bivalent domains |

Methylation of histone tails occurs mainly at lysine (K) and arginine (R) residues, most commonly observed as mono-, di-, or trimethylation of the lysine residues on H3 and H4 histone tails ^22–24^. Methylation of H3K36 and H3K4, for instance, act as activating histone modifications, while H3K27 methylation has a role in repressing the gene expression ^25–28^. Similarly, H3K27ac and H3K9ac modifications are associated with active transcription, and are predominantly associated with promoter and enhancer regions ^25–28^. Histone methyltransferase (HMT) are histone-modifying enzymes that catalyze the transfer of methyl groups to the targeted residues (lysine and arginine) through a domain known as SET domain ^29, 30^. Acetylation and deacetylation of histone tails, on the other hand, are catalyzed by histone acetyltransferases (HATs) and histone deacetylases (HDACs), respectively. Different chromatin-modifying enzymes, including histone deacetylases and histone lysine methyltransferases, function through multi-protein complexes that can also interact with methyl-CpG binding proteins, thereby linking mechanisms of histone modifications to the biochemical mechanism that maintains and modifies DNA methylation ^31, 32^. This implies that the enzymatic control of different epigenetic mechanisms are linked via crosstalk, and hence mutually interactive in regulating gene expression ^33^. In general, enzymes involved in depositing these chemical modifications (acetyl, methyl, etc.) onto the chromatin at the specific location are known as writers. In contrast, those which remove such modifications are called erasers ^34, 35^.

Despite the importance of the cnidarian Symbiodiniaceae relationship for ecosystem functioning we still know very little about the role of epigenetic mechanisms, and specifically histone modifications, in the regulation of host-symbiont interactions. Here, we profiled the genome-wide association of the histone modifications H3K27me3, H3K36me3 and H3K4me, H3K27ac and H3K9ac, in the cnidarian symbiosis model Exaiptasia diaphana (Aiptasia). We describe their genetic context, their correlation with CpG methylation (mCpG) and gene expression, as well as their association with, and putative regulation of symbiosis genes.

## Results

### Genome-wide distributions of histone modifications in *E. diaphana* and their correlation with CpG methylation

To understand the regulatory function of prominent histone modifications and their role in symbiosis, we performed ChIP-seq experiments in symbiotic *E. diaphana* and analyzed the genomic distribution of five major modifications; H3K27me3, H3K36me3, H3K27ac, H3K4me3 and H3K9ac (Supplementary Table ST1-ST5), as well as their correlations with respect to DNA methylation ^6^ and gene expression. To do this, we first called “peaks” from all ChIP-seq data, which are regions in the genome that are actively bound by a respective histone modification (see materials and methods for more details). We used the term “peak” throughout the manuscript to refer to these bound regions as well as their signal intensity relative to the input control.

For initial validation purposes, we compared the distribution of histone modifications between active and inactive regions of the *E. diaphana* genome ^36^. Repeat regions in the genome are mostly silenced ^37^ and are known to differ in bound histone modifications in comparison to non-repeat, i.e. genic, regions ^38^. Genome-wide analysis showed that the modifications H3K27me3 and H3K36me3 had significantly higher peaks (T-test; *p* < 0.0001) in repeat regions compared to genic regions. In contrast, H3K4me3, H3K27ac and H3K9ac had significantly higher peaks (T-test; *p* < 0.0001) in genic regions (Fig. 1A). This suggested that the transcriptional suppression of repeat elements in the *E. diaphana* genome aligns with higher H3K27me3 and H3K36me3 signals, and simultaneously lower peak signals for H3K4me3, H3K27ac and H3K9ac.

**Fig. 1:**
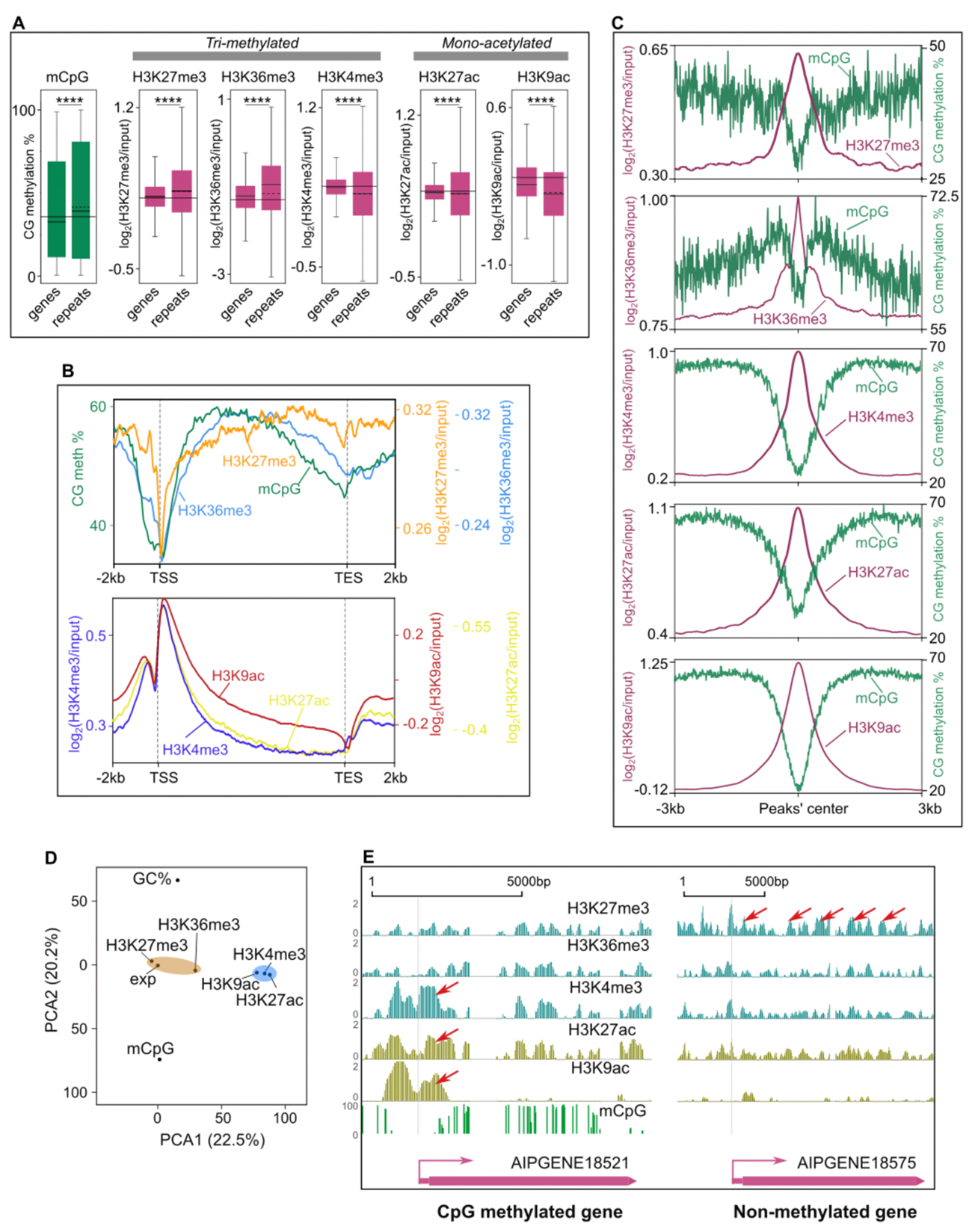
Genome-wide distribution of histone modifications in *E. diaphana* and their correlation with CpG methylation. **(A)** Boxplots of mCpG, H3K27me3, H3K36me3, H3K4me3, H3K27ac and H3K9ac peak levels in genes and repeats region of the *E. diaphana* genome. The solid horizontal line in each boxplot represents the median and the dotted line the mean. The solid horizontal line for each modification represents the average median of both genes and repeats (unpaired two-tailed Student’s t-test; \*\*\*\**p* < 0.0001). **(B)** Enrichment profiles of histone modifications and DNA methylation around all the protein-coding genes. x-axis is the gene locations from −2kb of TSS through gene-body and +2kb of TES; y-axis is the percentage of CpG methylation (mCpG) and log enrichment of peaks for each histone modification. **(C)** Average mCpG distribution around well-positioned histone modification’s peaks. **(D)** Principal-component analysis of all five histone modifications (H3K27me3, H3K36me3, H3K4me3, H3K27ac and H3K9ac), CpG methylation (mCpG), GC content (GC %), and gene expression (exp). **(E)** Genome browser snapshot showing example distribution of all five histone modifications (H3K27me3, H3K36me3, H3K4me3, H3K27ac and H3K9ac) in gene-body and promoter of methylated; AIPGENE18521 and non-methylated; AIPGENE18575 genes.

Next, we determined the peaks for each histone modification around all protein-coding genes in the *E. diaphana* genome. Peaks of H3K27me3 and H3K36me3 were prevalent in the gene-body and promoter regions, but not the transcriptional start site (TSS). In contrast, peaks of H3K4me3, H3K27ac and H3K9ac exhibited a bimodal peak pattern in the TSS region, with a smaller peak around the promoter region and a prominent peak around the first exon, while gene-bodies featured comparatively lower peaks (Fig. 1B, Supplementary Fig. S1A and S1B). To investigate the relationship between the different histone modifications and mCpG, we classified all genes from *E. diaphana* as either methylated (n=8018) or non-methylated (n=21322) based on their methylation density and methylation level (see materials and methods for more details). We found that H3K27me3 was predominantly present in the gene-body of non-methylated genes, while H3K36me3, H3K4me3, H3K27ac and H3K9ac were associated with methylated genes (Supplementary Fig. S1C – S1G, Supplementary Table ST6).

Interestingly, however, analysis of the core nucleosome regions of all five histone modifications showed either very low or no CpG methylation (Fig. 1C), suggesting that the DNA wound around the histone octamer core containing these modifications is mCpG depleted.

To determine how the different histone modifications are correlated with CpG methylation and gene expression, we analyzed their associations with mCpG, GC content and gene expression ^9^ using a principal-component analysis (Fig. 1D). All parameters, i.e., histone peak score, mCpG ratio, and GC content, were averaged for each gene. We observed that all histone modifications aligned on the same plane of the second principal component, along with gene expression. This suggests a tighter relationship between histone modifications and gene expressions compared to mCpG or GC content. Conversely, only histone modifications exhibiting gene-body prevalence aligned with mCpG, GC content and gene expression along the first principal component. This is likely because mCpG and GC content also display strong gene-body prevalence. Further, gene-body prevalent histone modifications (H3K27me3 and H3K36me3) and TSS prevalent modifications, (H3K4me3, H3K27ac and H3K9ac) clustered together, respectively (Fig. 1D). This suggests that the distribution patterns of mCpG and GC content are more similar to those of gene-body prevalent histone modifications. To further confirm this, we performed linear regression analyses to identify potential interactions between histone modifications at each gene (Supplementary Fig. S1H - S1K). We found a positive correlation between H3K4me3 and H3K9ac (*R*^2^ *= 0.27, p = 0.0023*), H3K27ac and H3K9ac (*R^2^ = 0.36, p = 0.0014*), and H3K27ac and H3K4me3 (*R^2^ = 0.43, p = 0.004*), which suggests that a substantial number of genes could potentially be bound by more than one of the three TSS dominated histone modifications (Supplementary Fig. S1H, S1I). In contrast, the correlation between gene-body-dominated histone modifications (H3K27me3 and H3K36me3) was very weak (*R^2^ = 0.052, p = 0.045*) (Supplementary Fig. S1J, S1K). Actual examples of the distribution of all five histone modifications over a methylated (AIPGENE18521) and non-methylated (AIPGENE18575) gene are shown in Fig. 1E.

### Histone-modification positioning and number as a function of transcription

Nucleosomes positioning and spacing has previously been shown to correlate with gene expression levels ^39^. To study this in *E. diaphana*, we analyzed how the positions of the different histone modifications in the promoter and gene-body regions vary and correlate with gene expression. To do this, we used the peak width as a parameter for position, with more precise nucleosome positioning reflecting tighter localization, and hence, narrower peak widths at a given position. For each modification, we compared the average peak width in the promoter and gene-body separately (Fig. 2A) across lowly, intermediately, and highly expressed genes. We found that the average peak width of activating histone modifications in the promoter region and the gene-body significantly increased with gene expression. Only the repressive modification H3K27me3 did not show a similar trend, and peak width remained constant across all expression levels.

**Fig. 2:**
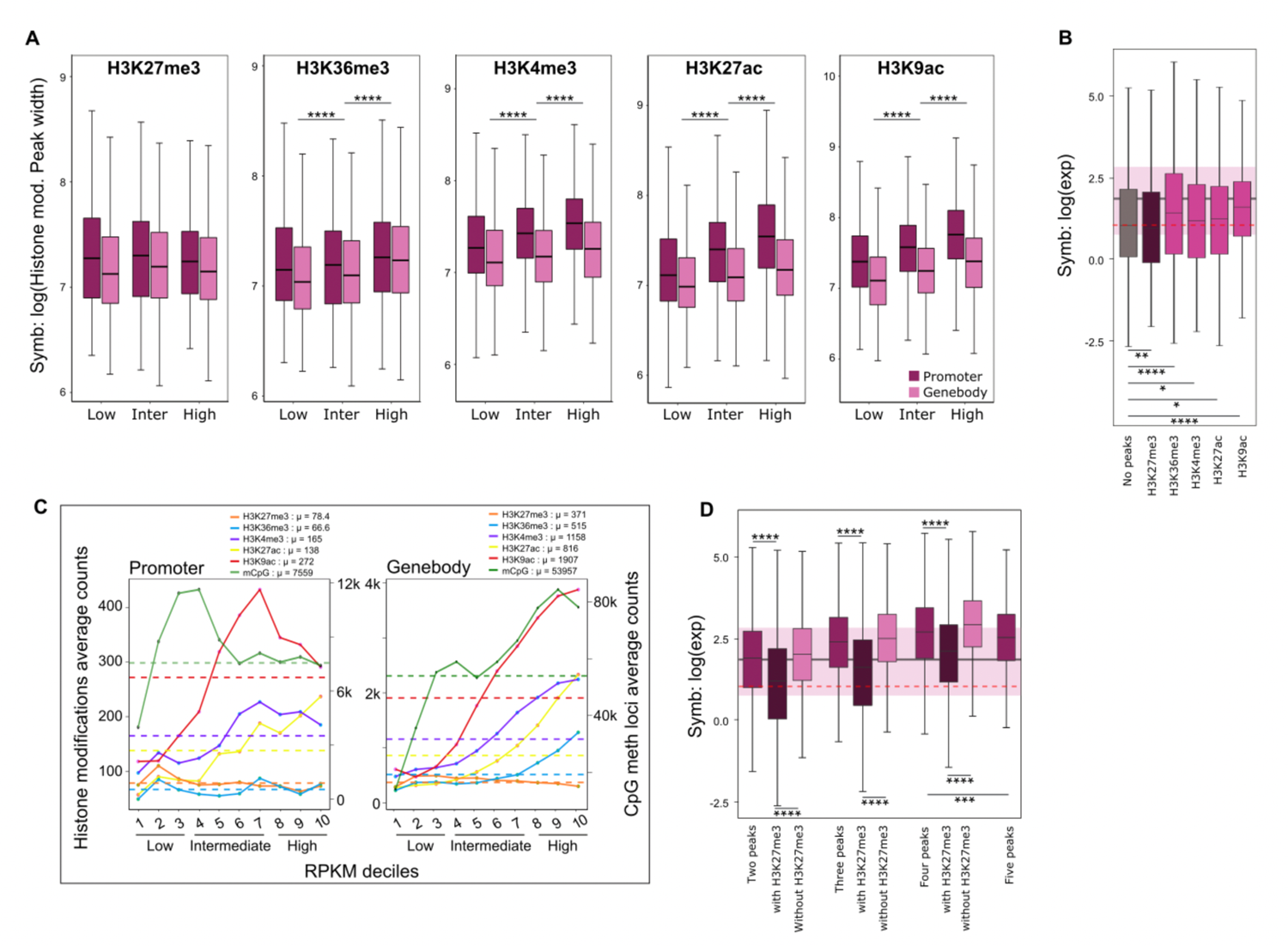
Histone-modification positioning and number as a function of transcription. **(A)** Average peak width of different histone modifications (H3K27me3, H3K36me3, H3K4me3, H3K27ac and H3K9ac) from promoter and gene-body as a function of gene expression category in symbiosis. Magenta boxplots are from promoters and pinks are from gene-bodies. X-axis is the gene category based on expression level (< 30^th^ percentile RPKM: low, 30^th^ and 70^th^ percentile RPKM: intermediate and > 70^th^ percentile RPKM: high); y-axis log value of peaks’ breadth (unpaired two-tailed Student’s t-test; \*\*\*\**p* < 0.0001). **(B)** Boxplot showing average expression for genes without peaks and genes with peaks for specific histone modifications. (unpaired two-tailed Student’s t-test; \**p* < 0.05, \*\**p* < 0.01, \*\*\**p* < 0.001, \*\*\*\**p* < 0.0001). **(C)** Specific histone modification counts are represented on the y-axis. X-axis represents gene expression values binned in deciles according to mRNA abundance (RPKM). Dashed lines represent the average count of each histone modifications and mCpG for all the genes, also shown in μ. Different chromatin modifications are represented by colors. **(D)** Boxplot showing average expression for the genes with two, three, four and five peaks. And correspondingly with and without the repressive modification H3K27me3; (unpaired two-tailed Student’s t-test; \*\*\**p* < 0.001, \*\*\*\**p* < 0.0001).

To investigate the effect of multiple peaks on gene expression, we selected genes with no peaks and compared their average expression level against genes with a single peak from one modification. We found that genes that only have a peak for the repressive modification (H3K27me3) exhibited a significantly lower average expression than genes without any peaks. At the same time, all genes with active modifications showed higher gene expression values than genes without any peak or with H3K27me3 (Fig. 2B). Furthermore, we found a general increment in gene expression with an increasing number of peaks for active histone modifications. Meanwhile repressive histone modification peak counts, H3K27me3, showed a negative and weak relation with expression (Fig. 2C). We also examined the cumulative effect of multiple histone Modifications on genes expression. Interestingly, we observed that the average gene expression level was higher if genes were associated with more than one histone modification, with every additional modification resulting in significantly higher expression levels of associated genes, as long as the repressive modification H3K27me3 was not included (Fig. 2D). Inclusion of H3K27me3 consistently correlated with significantly lower gene expression levels, further confirming its repressive effect.

### Histone modifications in *E. diaphana* correlate with gene expression

To further analyze the correlation of the different histone modifications with mCpG and gene expression, we divided all the protein-coding genes from *E. diaphana* into six categories based on their transcription level and DNA methylation status. We observed a positive correlation between histone peak height and gene expression for H3K36me3 and for all TSS-prevalent histone modification (H3K4me3, H3K27ac, and H3K9ac), and only H3K27me3 displayed a negative correlation (Fig. 3A). This finding affirmed our previous analysis showing that H3K27me3 peak counts decreased with increasing gene expression (Fig. 2C).

**Fig. 3:**
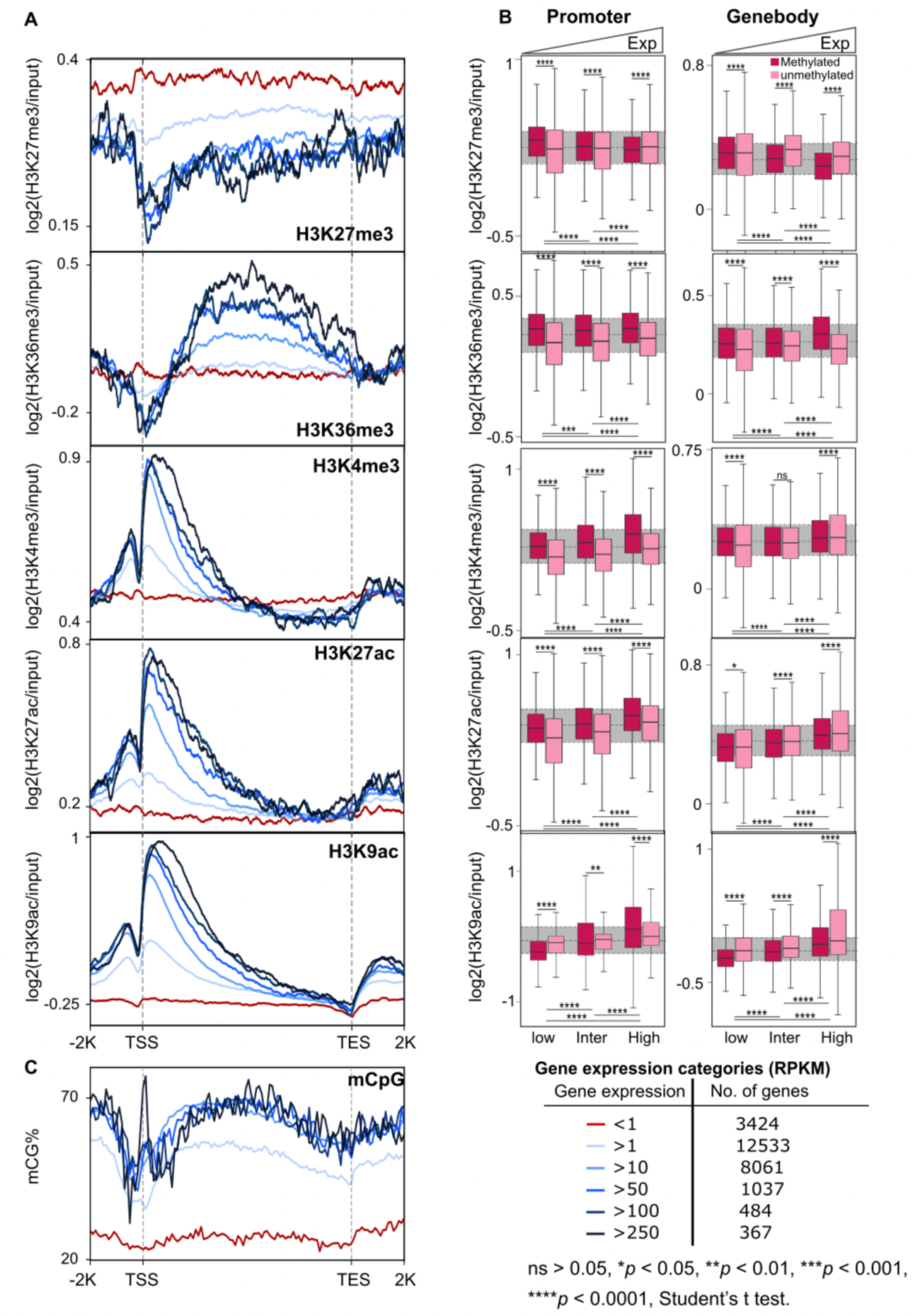
Histone modifications in *E. diaphana* correlate with gene expression. **(A)** Distribution of histone modifications around genes with increasing expression levels. Red line marking the lowest gene expression category (RPKM < 1), and darkest blue the highest expression category (RPKM >250). **(B)** Boxplots of histone peak heights from promoter and gene-body regions of methylated (dark pink) and unmethylated (light pink) genes. Genes with expression below the 30^th^ percentile of RPKM were classified as lowly expressed, those between the 30^th^ and 70^th^ percentile as intermediately expressed, and those above the 70^th^ percentile as highly expressed. (unpaired two-tailed Student’s t-test; ∗∗*p* < 0.01, ∗∗∗*p* < 0.001, ∗∗∗∗*p* < 0.0001). **(C)** mCpG distribution around genes with increasing gene expression levels.

To further investigate the potential interactions of histone modifications and mCpG, we plotted the average peak heights for every histone modification for low, intermediate and highly expressed genes, each with and without mCpG respectively (Fig. 3B). We found that H3K27me3 showed a significant negative correlation (*p* < 2.2 × 10^-16^) with gene expression both when present in the promoter or the gene-body, and this effect was even more pronounced in methylated genes. In contrast, H3K36me3 showed a positive correlation with mCpG and gene expression (Fig. 3B), with H3K36me3 peak height positively correlating with increasing expression in methylated genes. Similarly, TSS-prevalent histone modifications, i.e., H3K4me3, H3K27ac and H3K9ac, also showed a positive correlation with gene expression and methylation (*p* < 2.2 × 10^-16^), and this effect was more pronounced for peaks in the promoter region (Fig. 3A and 3B).

### Histone modifications regulate the transcriptional response to symbiosis

To investigate the role of histone modifications in the regulation of the symbiotic relationship between *E. diaphana* and its dinoflagellate symbionts, we analyzed the correlation between peak occupancy for each histone modification and gene expression across 731 previously identified symbiosis-associated genes ^9^. We first categorized these 731 symbiosis-associated genes into symbiosis-repressed (365) and symbiosis-induced (366) genes. We found that most symbiosis genes (544; 74.4% of the 731 genes) were associated with at least one of the five histone modifications we analyzed (Fig. 4A, Table 2, Supplementary Table ST7 – ST11). Active histone modifications (i.e., H3K36me3, H3K4me3, H3K27ac and H3K9ac) showed a significantly higher association with symbiosis-induced genes (*p < 0.01*) while the repressive histone modification (H3K27me3) had an almost equal number of peaks in both categories.

**Fig. 4:**
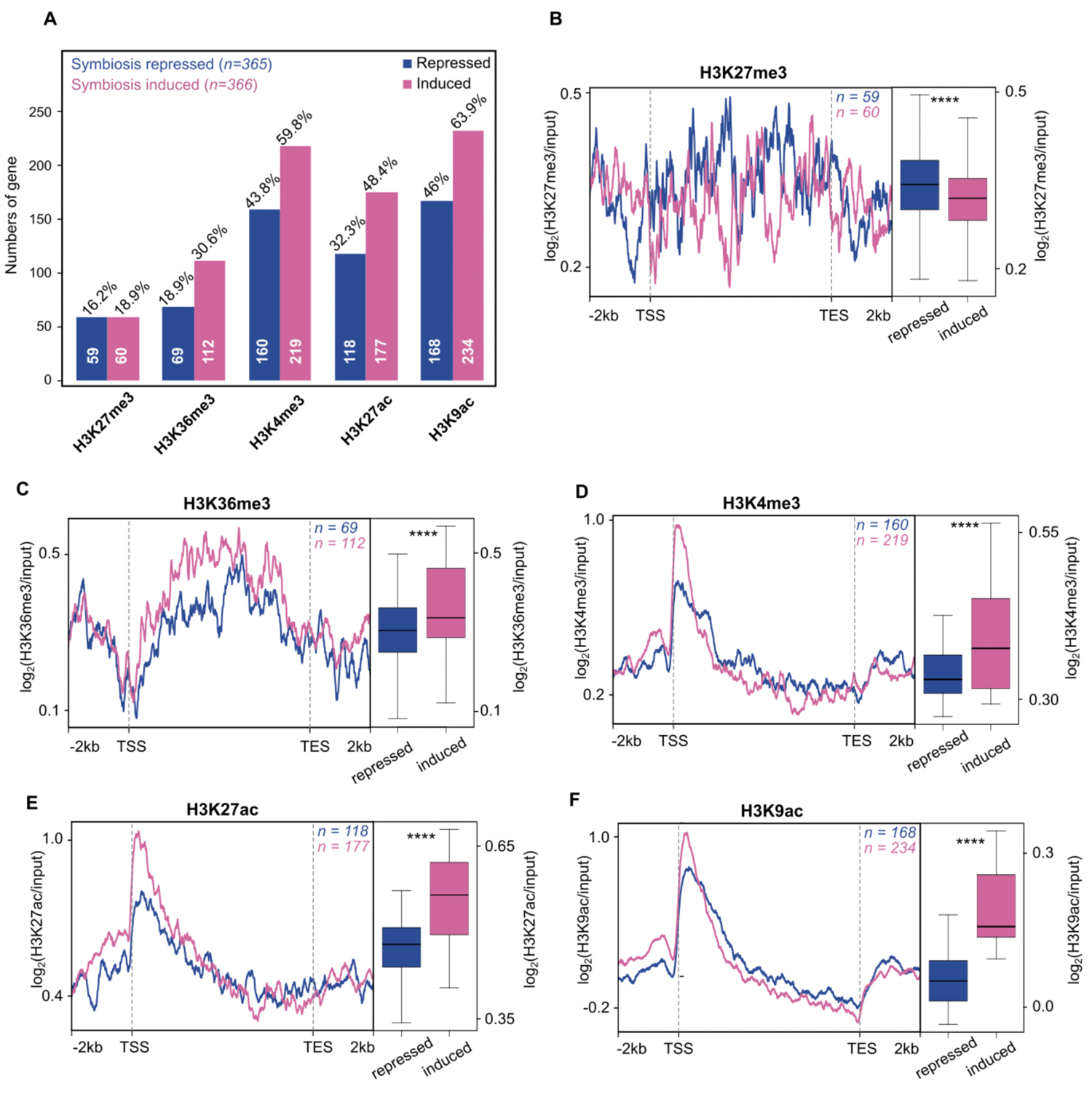
Histone modifications regulate the transcriptional response to symbiosis. **(A)** Total number of symbiosis-repressed (blue) and induced (pink) genes associated with each histone modification in their gene-body and promoter regions. Average peak distributions of symbiosis-repressed (blue) and induced (pink) genes associated with H3K27me3 **(B)**, H3K36me3 **(C)**, H3K4me3 **(D)**, H3K27ac **(E)** and H3K9ac **(F)** from −2kb of TSS through gene-body and +2kb of TES. Each of the line plots from symbiosis-repressed genes is compared with induced genes (see respective boxplots, unpaired two-tailed Student’s t-test; ****p < 2.2 × 10^-16^).

**Table 2:** Genes associated with histone modifications in the *E. diaphana* genome and symbiosis genes, respectively

| Histone modifications | H3K27me3 | H3K36me3 | H3K4me3 | H3K27ac | H3K9ac | # of genes with at least one modification |
| --- | --- | --- | --- | --- | --- | --- |
| **Total in genome** | 5660 | 6536 | 13664 | 9629 | 14738 | 18675 (63.8%) |
| **Symbiosis-associated genes** | 119 (16.3%) | 181 (24.8%) | 379 (51.8%) | 295 (40.4%) | 402 (55%) | 544 (74.4%) |
| **Symbiosis-repressed genes** | 59 (16.2%) | 69 (18.9%) | 160 (43.8%) | 118 (32.3%) | 168 (46%) | 252 (69%) |
| **Symbiosis-induced genes** | 60 (18.9%) | 112 (30.6%) | 219 (59.8%) | 177 (48.4%) | 234 (63.9%) | 292 (79.8%) |

To confirm the relationship between peak height and gene expression, we compared the average histone peak height of every modification across symbiosis repressed and induced genes. We found that the repressive modification H3K27me3 had significantly higher peaks in symbiosis-repressed genes (*p <* 2.2 × 10^-16^, Fig. 4B), while all active modifications (H3K36me3, H3K4me3, H3K27ac and H3K9ac) had significantly higher peaks (*p <* 2.2 × 10^-16^) in symbiosis-induced genes (Fig. 4C – 4F). This finding confirmed that histone modifications play an active role in the regulation of the symbiotic relationship between *E. diaphana* and its dinoflagellate symbiont.

As a final step of validation, we analyzed the correlation between histone modifications and changes in gene expression in response to symbiosis, we divided both symbiosis-induced (n=366) and symbiosis-repressed genes (n=365) into two groups based on their median gene expression fold change. We compared the profiles of the upper and lower 50^th^ percentile for each of the histone modifications separately. Similar to our previous observation we found that the repressive modification H3K27me3 showed a higher prevalence in symbiosis-repressed genes irrespective of the fold change (*p <* 2.2 × 10^-16^, Supplementary Fig. S2A). Conversely, our analysis on the active histone modifications; H3K36me3, H3K4me3, H3K27ac and H3K9ac, also confirmed our previous findings of significantly higher peaks in symbiosis-induced genes (*p* < 2.2 × 10^-16^) (Supplementary Fig. S2B – S2E).

### Histone modifications are involved in symbiosis-induced nutrient metabolism

To understand the role of histone modifications in the regulation of the symbiotic relationship, we performed gene ontology (GO) enrichment analyses for both symbiosis repressed and induced genes for each histone modification individually (Supplementary Table ST12 – ST21, Supplementary Fig. S3). While each modification had several unique enriched biological functions, we found a considerable number of categories that were enriched across two or more histone modifications (Supplementary Table ST22 – ST23). Many of these shared categories within the symbiosis-induced genes were involved in amino acid metabolic processes, such as the regulation of cellular amino acid and protein metabolic process (Table 3). Furthermore, we found processes involved in the response to glucose and ammonium transport to be associated with multiple modifications, suggesting that the central function of this symbiotic relationship in driving host amino acid biosynthesis is regulated through histone modifications ^9^. In line with this, we also found shared enrichment of amino acid biosynthesis related processes, including L-serine, glutamine, and methionine, as well as categories involved in amino acid transport. Similarly, we found several shared biological processes associated with organism growth to be enriched across multiple histone modifications, most notably GO terms related to central growth pathways such as the insulin, hippo and the TORC1 pathways.

**Table 3:** Selected GO terms of symbiosis genes associated with histone modifications. The full GO term list is shown in Supplementary Tables (ST12 – ST23).

| GO Term | Description |
| --- | --- |
| GO:0000098 | sulfur amino acid catabolic process |
| GO:0003333 | amino acid transmembrane transport |
| GO:0006521 | regulation of cellular amino acid metabolic process |
| GO:0006527 | arginine catabolic process |
| GO:0006541 | glutamine metabolic process |
| GO:0006559 | L-phenylalanine catabolic process |
| GO:0006564 | L-serine biosynthesis process |
| GO:0006579 | amino-acid betaine catabolic process |
| GO:0009086 | methionine biosynthesis process |
| GO:0009115 | xanthine catabolic process |
| GO:0009749 | response to glucose stimulus |
| GO:0015804 | neutral amino acid transport |
| GO:0019518 | L-threonine catabolic process to glycine |
| GO:0031931 | TORC 1 complex |
| GO:0032024 | positive regulation of insulin secretion |
| GO:0035329 | hippo signaling pathway |
| GO:0044267 | cellular protein metabolic process |
| GO:0046168 | glycerol-3-phosphate catabolic process |
| GO:0072488 | ammonium transport |
| GO:0046949 | fatty-acyl-CoA biosynthetic process |
| GO:0046500 | S-adenosylmethionine metabolic process |
| GO:0006556 | S-adenosylmethionine biosynthetic process |
| GO:0008898 | S-adenosylmethionine-homocysteine S-methyltransferase activity |
| GO:0032259 | methylation |
| GO:0001733 | galactosylceramide sulfotransferase activity |
| GO:0003943 | N-acetylgalactosamine-4-sulfatase activity |
| GO:0003810 | protein-glutamine gamma-glutamyltransferase activity |
| GO:0070403 | NAD<sup>+</sup> binding |
| GO:0004029 | aldehyde dehydrogenase (NAD<sup>+</sup>) activity |

While many of the enriched categories in the symbiosis-induced genes were associated with anabolic processes, we found symbiosis-repressed genes associated with histone modifications to be predominantly involved in catabolic processes. These included the catabolism of molecules like xanthine and glycerol-3-phosphate but also amino acids such as sulfur amino acids, L-phenylalanine, betaine, arginine and L-threonine, among others.

Interestingly, we also found enrichment of GO terms involved in various metabolic pathways that generate metabolites important for epigenetic modifications, such as acetyl-CoA/fatty-acyl-CoA, *S*-adenosylmethionine, methylation process and lactate. Previous studies have shown that these metabolites serve as cofactors for the enzymes responsible for depositing the chemical modifications (acetyl and methyl) onto chromatin; chromatin writers ^34,35^. In addition, we found GO terms related to metabolites such as α-ketoglutarate and NAD+, which are essential cofactors for certain enzymes that remove chemical modifications; chromatin erasers ^34, 35^. This suggests that the chromatin changes induced to regulate gene expression in response to symbiosis might be supplied by these processes and ultimately established through the respective writers and erasers.

## Discussion

The process of symbiosis establishment and maintenance requires changes in the cnidarian host’s cell function and specialization. Epigenetic mechanisms have been shown to play critical roles in symbiotic relationships of eukaryotic and bacterial cells ^16^. The general importance of histone modifications in host-microbe interactions has been acknowledged in plants, humans, and other invertebrates ^16–18^. Through chemical signals and metabolites, endosymbionts can influence epigenomes of host cells and directly enable communication between the two partners ^18, 40^. Interestingly, histone acetylase and deacetylase activity have been shown to be influenced by microbes and dietary factors ^15, 41, 42^. Although epigenetic studies in cnidarians remain scarce, there is evidence that histone modifications may play a critical role in host-algae symbiosis mechanisms ^6, 43, 74, 75^. Here, we report the first genomic landscape of five histone modifications, H3K27me3, H3K36me3, H3K4me3, H3K27ac and H3K9ac, in a symbiotic cnidarian.

We find that their genomic distribution and putative primary functions align with observations made in other organisms ^44^, suggesting functional conservation of these histone modifications in *E. diaphana*. Further, our results revealed strong correlations between the histone modifications analyzed and transcriptional changes observed in response symbiosis. These findings collectively suggest a direct role for histone modifications in regulating the host’s response to symbiosis.

### Conserved roles of histone modifications in regulating gene expression

The general explanation for the ability of histone modifications to enhance or repress transcription is that they affect the DNA-histone association and, thus, promote or suppress access for transcription factors and the transcriptional machinery to the DNA. As such, these modifications represent an essential mechanism for the epigenetic control of transcriptional responses in eukaryotes ^12, 45–47^. In line with this, our analyses revealed highly significant correlations between histone modifications and gene expression. Analysis of the genomic distribution of H3K27me3 and H3K36me3 showed enrichment in repeat regions ^45, 46^ while the activating modifications H3K4me3, H3K27ac and H3K9ac showed enrichment in the genic regions (Fig. 1A), as expected based on observations in other organisms ^11, 48^.

It is interesting to note, however, that we found a substantial number of genes associated with more than one histone modification, suggesting that several histone modifications might act on the same gene simultaneously (Supplementary Fig. S1H – S1K). Such a cooperative interaction in regulating gene expression was further supported by the finding that the number of active histone modifications present on genes positively correlated with gene expression levels, suggesting an additive effect. However, it needs to be pointed out that ChIP-seq data cannot inform if the modifications were present on the same DNA molecule or if they were just associated with the same gene but in different cells of the organism. This limitation is evident when looking at the lower average expression observed for genes that were bound by an activating histone modification as well as the repressive modification H3K27me3. Since an actively expressed gene is unlikely to be simultaneously associated with activating and repressive modification, it is more likely that the H3K27me3 association stems from cells where this gene was silenced. Since these cells would not contribute any transcripts for this gene to the whole organism RNA pool, this would reduce the observed overall expression level of the gene in the organism.

### Crosstalk between histone code and DNA methylation

DNA has a determined nucleotide sequence that cannot be changed. However, it has been postulated that the transcription of the genetic information is partly regulated by epigenetic mechanisms such as the underlying histone modifications and DNA methylation. Our analyses of potential interactions of histone modifications and DNA methylation in regulating gene expression revealed strong correlations for all activating modifications that suggest crosstalk between these epigenetic mechanisms in *E. diaphana*. We observed that the average expression of genes associated with activating histone modifications was generally higher if they were also methylated (Fig. 3B), suggesting a cooperative interaction between activating histone modifications and DNA methylation. In contrast, we found that genes associated with the repressive modification H3K27me3 showed the opposite trend for methylated genes if the histone modification was found in the gene-body. However, it should be noted that H3K27me3 was predominantly present in the gene-body of non-methylated genes, while H3K36me3, H3K4me3, H3K27ac and H3K9ac were associated with methylated genes (Supplementary Fig. S1C – S1G). While these results suggested a cooperative interaction, analysis of the core nucleosome regions of all five histone modifications showed either very low or no CpG methylation (Fig. 1C), indicating that they are present on the same genes but that their precise locations within the gene are mutually exclusive. In summary, our results are indicative of crosstalk between active histone modifications and DNA methylation in modulating gene expression, while repressive modifications associate predominantly with non-methylated genes to suppress their expression.

### A model for the regulation of gene expression via histone modifications in *E. diaphana*

Based on our results, we propose a model for how the histone modifications analyzed here could regulate gene expression in *E. diaphana*. In addition, the model demonstrates how histone modification and DNA methylation crosstalk may be functioning in symbiotic cnidarians.

When a gene is silenced, it is bound by H3K27me3 in the promoter and gene-body (Fig. 5A), which promotes repression through the polycomb complex. H3K27me3 has been shown to recruit PRC1 (polycomb repressive complex), which contributes to the compaction of the chromatin, leading to the formation of heterochromatin and its inaccessibility for transcription factors and the transcriptional machinery ^49^.

**Fig. 5:**
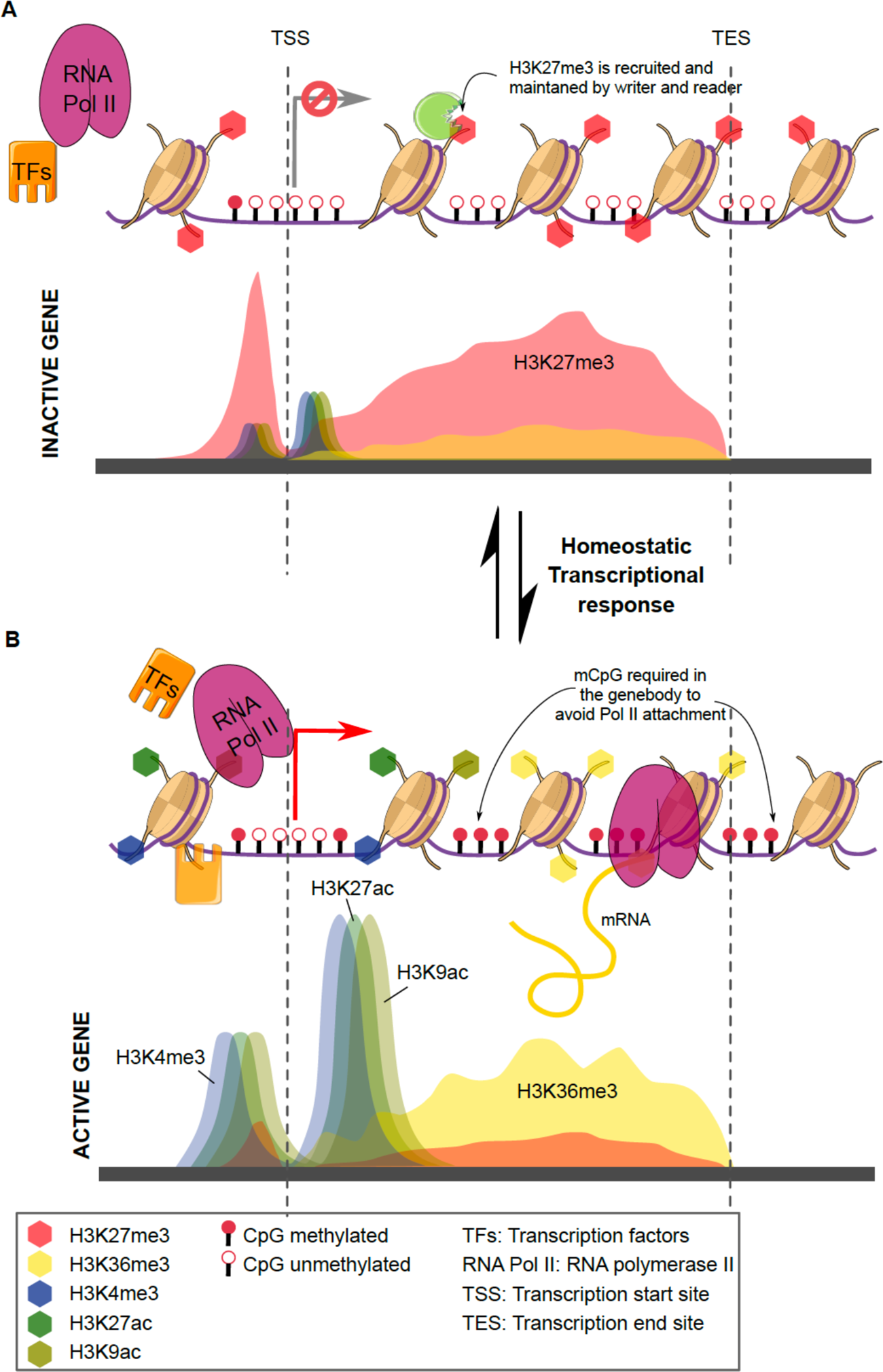
Histone modifications and CpG methylation underlying dynamic gene regulation. Proposed model for the dynamic topology of five histone modifications and CpG methylation on gene loci: (**A)** Chromatin erasers remove all active histone modifications and DNA methylation from the gene-body (H3K36me3 and CpG) and promoter (H3K4me3, H3K27ac, H3K9ac and CpG) of an inactivated gene. Simultaneously, chromatin writers add the repressive mark H3K27me3 to histone H3 molecules within the promoter and gene-body. This chromatin state prevents RNA Pol II and transcription factors (TFs) from attaching to the promoter. **(B)** For the activation of gene expression H3K4me3, H3K27ac and H3K9ac are established at the promoter and the transcriptional start sites of the gene, while H3K36me3 and CpG methylation are established throughout the gene-body. This facilitates access of Pol II and TFs to the promoter and the TSS, which activates the gene and promotes transcription.

However, when a gene needs to be activated, it requires the preinitiation complex (PIC) to assemble at the promoter region of a gene and to recruit RNA Pol II to the promoter to build the transcription initiation complex ^50–52^. Based on our results, and in line with previous findings ^11, 27, 53^, we propose that the presence of the activating histone modifications H3K27ac and H3K9ac around the promoter and the TSS promote access for transcription factors of the PIC to the gene promoter (Fig. 5B). Once the PIC is assembled, RNA Pol II can be recruited to form the transcription initiation complex and H3K27ac, H3K9ac and H3K4me3 (TSS-dominated modifications) can act as a pause-release signal for Pol II to initiate transcription. The transcriptional elongation process is then supported by H3K4me3 and H3K36me3 ^54^. Meanwhile, low CpG methylation at the promoter further favors the attachment of the assembly of the PIC and the transcriptional complex ^55^, while high CpG methylation in the gene-body prevents the assembly of the transcriptional machinery at cryptic promoter sequences within the gene-body, which would lead to spurious transcripts and the production of truncated proteins ^56^. The crosstalk between histone modifications and DNA methylation is brought about through the interaction of histone modifying enzymes. For instance, the histone methyltransferase Set2D is recruited by the active transcriptional complex and tri-methylates H3K36 along the gene-body. H3K36me3, in turn, is then actively bound by DNA methyltransferase 3b which methylated CpG within the gene-body of actively transcribed genes ^57^. Together, histone modifications and DNA methylation create a chromatin landscape conducive of high gene expression and the production of full-length transcripts, while at the same time reducing transcriptional noise and spurious transcripts (Fig. 5B)^58^.

### The role of histone modifications in symbiosis

Our analyses revealed that genes associated with activating histone modifications were significantly enriched in the fraction of symbiosis-induced genes. This suggests that their increased expression in response to symbiosis is promoted via their association with activating histone modifications. Interestingly, we did not see the opposite trend for the repressive modification H3K27me3 which is associated with the same number of symbioses induced and repressed genes. However, analysis of H3K27me3 peak heights did show significantly higher peaks in symbiosis-repressed genes compared to symbiosis-induced ones. The fact that H3K27me3 peaks were significantly higher in symbiosis-repressed genes suggests that the association of these genes with H3K27me3 was evident in more host cells, which increased the number of ChIP-seq reads obtained, and thus the relative peak heights.

Analyses of the biological functions enriched in 2 or more histone modifications highlighted that the histone modifications studied associated with anabolic functions in symbiosis-induced genes and catabolic functions in symbiosis-repressed genes. Specifically processes associated with amino acid biosynthesis and growth suggested that these histone modifications are involved in regulating the metabolic response and growth in symbiotic anemones. However, these processes are also involved in the regulation of the symbiotic relationship itself. Endosymbiotic relationships are usually driven by synergies arising from the complementation of the host’s metabolic capabilities that enable the resulting metaorganism to thrive in nutrient poor environments or to use previously inaccessible diets ^59, 60^, as is also the case for symbiotic anemones and corals. However, this intimate form of symbiosis requires maintaining a delicate balance of nutrient fluxes to provide nutrients to the symbionts, to keep benefiting from them, but at the same time ensure they do not over proliferate at the expense of the host. The maintenance of this balance in symbiotic cnidarians is achieved through the regulation of genes involved in ammonium assimilation and amino acid biosynthesis ^9^. We find that genes involved in the assimilation of waste ammonium and amino acid biosynthesis are predominantly associated with activating histone modifications. Further, the observed crosstalk of activating histone modifications and DNA methylation in driving higher expression of symbiosis-induced genes suggest a multi-layer epigenetic regulatory mechanism that may be critical for cnidarian symbiosis. Our results, therefore, indicate that the maintenance of symbiosis-associated gene expression is provided by the synchronous action of histone modifications and DNA methylation. However, the biological interpretation of the results presented here are only first insights into research that clearly requires further expansion. We acknowledge that to further disseminate the role of histone modifications in symbiosis, further ChIP-seq studies including aposymbiotic individuals (*E. diaphana* without its dinoflagellate symbionts) will be required. Nonetheless, our results on the biological functions align with recent observations of increased gene accessibility and open chromatin states in symbiotic anemones ^43^.

## Materials and Methods

### *E. diaphana* culture and maintenance

*E. diaphana* of the clonal strain CC7 ^61^, originating from North Carolina, was used in this study. Anemones were maintained in polycarbonate tubs with autoclaved seawater at 25°C. Animals were exposed for 12-hour light/dark cycle at 20-40 μmol photons m^2^s^1^ light intensity. The anemones were fed twice weekly with freshly hatched *Artemia nauplii* (brine shrimp). For the experiment, three biological replicates were taken, each consisting of two individual anemones pooled together.

### Chromatin Immunoprecipitation (ChIP) sequencing library preparation

The process of establishing a reproducible ChIP-seq protocol in *E. diaphana*, which so far has primarily been optimized for human, mice and plant cell studies, included many quality control and optimization steps that require attention. In hopes of streamlining future attempts at ChIP-seq in other cnidarians, especially corals, we opted to optimize pre-immunoprecipitation steps to the point that kits could be confidently used thereafter. We used Zymo-Spin ChIP Kit (Zymo Research) to extract histone-bound DNA fragments, however, we applied minor adjustments to the pre-IP steps. In a recent publication ^6^, we published a summarized version of the protocol. In another publication ^62^, a summarized version of the protocol was presented. Here, we provide a detailed description of the protocol (Supplementary file – Extended materials); steps of validation and optimization are described in greater detail to, hopefully, allow future research to progress and further advance the field of epigenetic research in cnidarians.

Corresponding input controls for each of the three replicates were generated as suggested by the manufacturer. After validation of various histone antibodies (Supplementary file – Extended materials; Fig. S3), immunoprecipitation was conducted using a target-specific antibody to histone 3 acetylation at lysine 27 – H3K27ac (ab4729, Abcam), histone 3 tri-methylation at lysine 4 – H3K4me3 (ab8580, Abcam), histone 3 acetylation at lysine 9 – H3K9ac (ab10812, Abcam), histone 3 tri-methylation at lysine 36 – H3K36me3 (ab9050, Abcam) and histone 3 tri-methylation at lysine 27 – H3K27me3 (ab6002, Abcam). Upon validation of immunoprecipitation, using High Sensitivity DNA Reagents (Agilent Technologies, California, United States) on a Bioanalyzer, ChIP libraries were constructed using TruSeq Nano HT DNA kit (Illumina, California, United States).

### Sequencing libraries

Paired and single end sequencing was performed at the Bioscience Core Lab (BLC) at the King Abdullah University of Science and Technology, Thuwal, KSA with NextSeq 500. The ChIP-seq mapped files are deposited in NCBI SRA under accession number PRJNA826667.

### Sequence alignments

ChIP-seq sequencing resulted in 10 million read pairs per replicate. The raw reads’ quality were checked with FASTQC toolkit ^63^ and cleaned to achieve desired quality using the and Trimmomatic ^64^. The clean reads were uniquely mapped on the *E. diaphana* genome (http://Exaiptasiadiaphana.reefgenomics.org/) ^36^ using bowtie 1.1.2 with default parameters ^65^.

### Identification and annotation of histone modification peaks

Genomic regions having all five modification modifications were identified using Model-based Analysis of ChIP-Seq (MACS3: https://github.com/macs3-project/MACS/tree/master/MACS3) through “*macs3 callpeak -t treatment.bam -i input.bam -f BAM -g 2.7e+8 -B --nomodel --d-min 10 --call-summits”* parameters ^66^. Combined evidence from ChIP-seq biological replicates were estimated by MSPC tool (https://github.com/Genometric/MSPC) ^67^, using *“./mspc -I rep*.bed -r bio -w 1e-4 -s 1e-8”* parameter. Absolute enrichments were calculated as log2(average signal/average input control) with adjusted *p* < 0.01 for each identified peak in whole genome.

We used gene annotation of the *E. diaphana* genome (GFF3 file) ^36^ for assigning the location of identified histone enrichment peaks on the genome. For annotation of all such peaks, we used ChIPseeker: An R/Bioconductor package for ChIP peak annotation, comparison and visualization^68^. This package annotates the peaks into the genic or intergenic region, and the distances to the 5′ and 3′ ends of each genomic feature (gene/intergenic region/promoter/exon/intron). The annotated table of all histone enrichments peaks are shown in Supplementary Tables ST1–ST5.

### Defining biological function of histone peaks

Gene annotation was obtained from the previously published *E. diaphana* genome ^36^. To analyze the functional enrichment of the histone peaks bound genes we obtained the GO annotation from the genome and analyzed it using topGO ^69^ using default settings. In order to test for the potential role of all five histone modifications in symbiosis, we used the list of 731 identified symbiosis genes as identified in *Cui et al., 2019* ^9^ and matched them with binding sites of each histone modification.

### Use of previously published data

DNA methylation BS-seq raw reads were obtained and processed from *Li et al., 2018* ^6^, and gene expression data were taken from *Cui et al., 2019* ^9^. Based on *Cui et al., 2019*, classification of symbiotic-dependent and -independent genes has been done.

### Data visualization

Screenshots of *E. diaphana* chromosome features were taken in Integrated Genome Browser-9.1.8 (IGB) ^70^. Previously used DNA methylation BS-seq data were processed with Bismark-0.22.3 ^71^ for each replicate, and then taken an average of replicates from each sample using basic Unix commands. Average enrichment scores, plots and heatmaps at genomic features of interest were generated by deepTools ^72^. Histone width and count for each associated gene of specific modification were estimated and plotted by in-house programs.

### Quantification and statistical analysis

All the other plots and boxplots with statistical analyses have been done mainly by R CRAN package: dplyr. Further statistical analyses were done using R (version 3.5.1). For boxplots, the bottom and top of the box indicate the 25^th^ and 75^th^ percentile, respectively. The bar in the boxplot shows the median. Whiskers indicate a 1.5X interquartile range (IQR).

### Data access

The ChIP-seq mapped files generated for this study have been submitted to the NCBI Gene Expression Omnibus (http://www.ncbi.nlm.nih.gov/sra/) under accession number PRJNA82666793.

## Author Contributions

M. A. and M. J. C. designed the project; M. J. C. performed the experiments; K.N., M.J.C, G.C. and K. G. M. analyzed the data; and K.N., M. J. C., and M. A. wrote the paper.

## Classification

Biological Sciences; Climate adaptation.

## Supplementary Figures

**Fig. S1:**
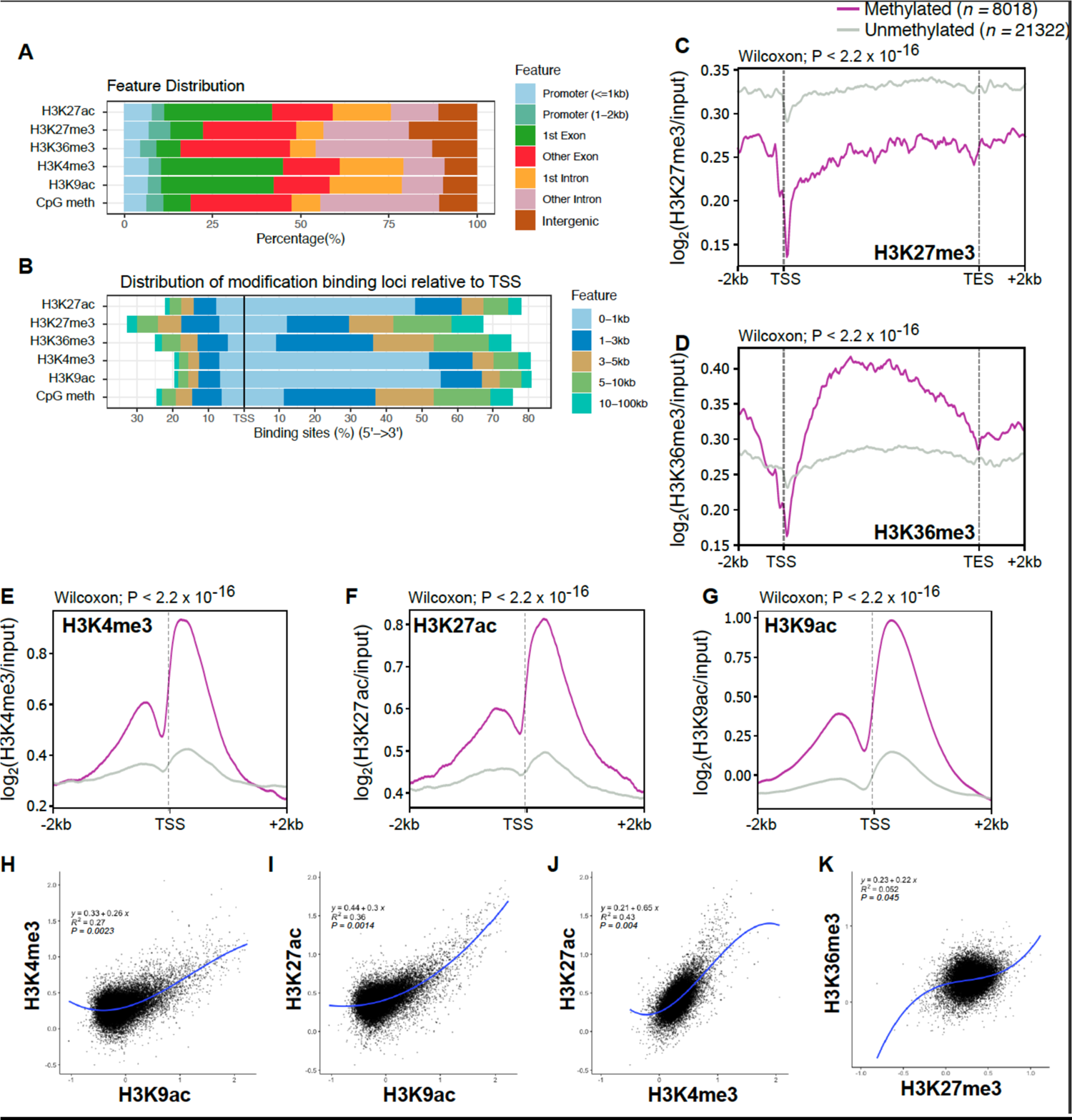
Genome-wide histone modifications distributions in *E. diaphana* and their correlations. **(A)** Genomic distribution of significantly identified peaks from different histone modifications and mCpG in Aiptasia genome. **(B)** Distribution of significantly identified peaks from different histone modifications and mCpG with respect to TSS in Aiptasia genome. Average peaks of methylated (pink) and unmethylated (grey) genes associated with H3K27me3 **(C)**, H3K36me3 **(D)**, H3K4me3 **(E)**, H3K27ac **(F)** and H3K9ac **(G)** from −2kb of TSS through gene-body and +2kb of TES. Linear regression analyses between H3K9ac with H3K4me3 **(H)**, H3K9ac with H3K27ac **(I)**, H3K4me3 with H3K27ac **(J)**, and H3K27me3 with H3K36me3 **(K)**.

**Fig. S2:**
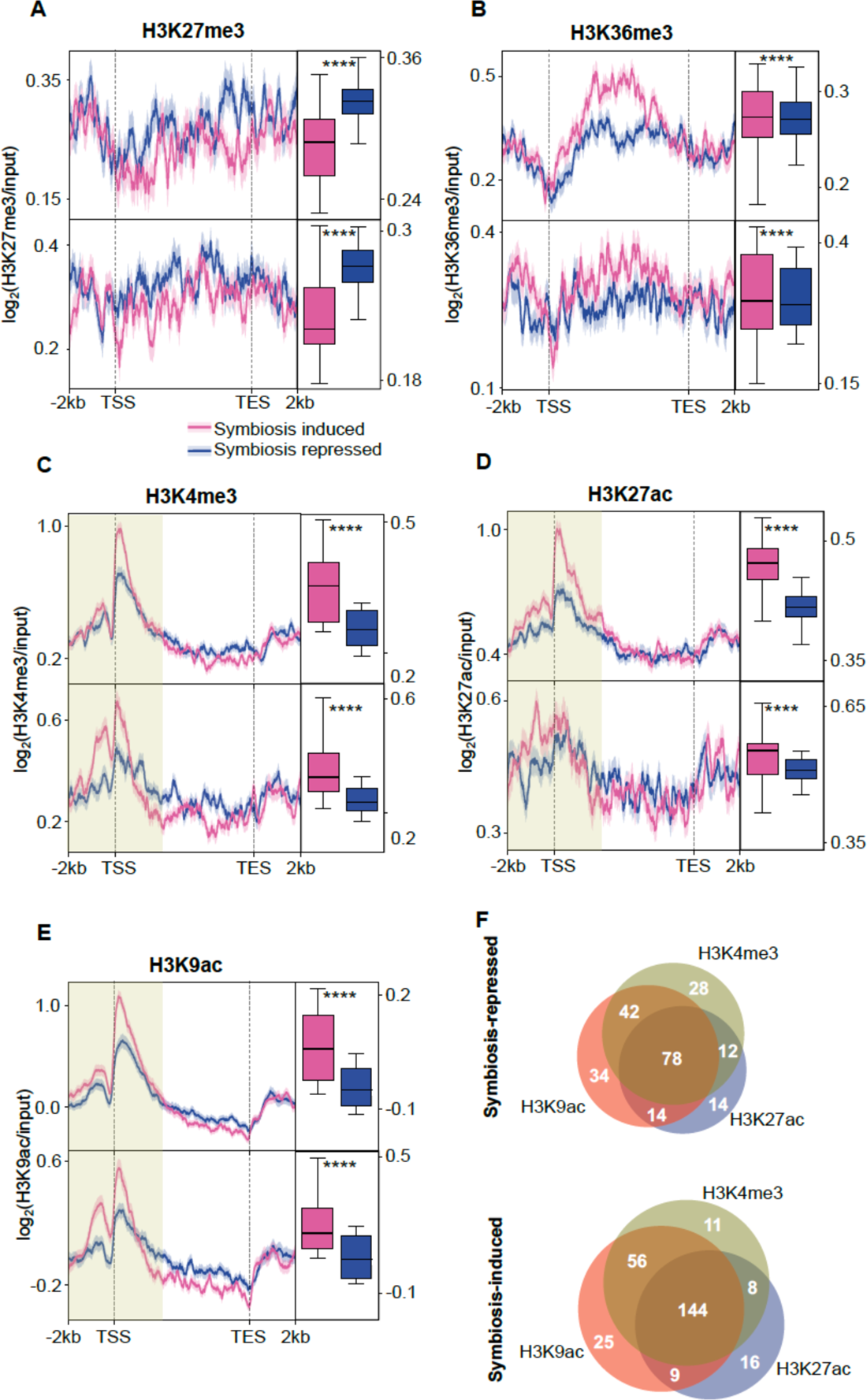
Upper and lower 50% percentile: histone modifications change within symbiosis induced and repressed genes. Symbiosis-repressed (n=365) and symbiosis-induced genes (n=366) genes were sub-divided into two groups based on their median expression fold change into upper and lower percentile of gene expression and their histone signals in H3K27me3 **(A)**, HeH36me3 **(B)**, H3K4me3 **(C)**, H3K27ac **(D)** and H3K9ac **(E)**. **(F)** Shared TSS dominated histone peaks in symbiosis-repressed and induced genes.

**Fig. S3:**
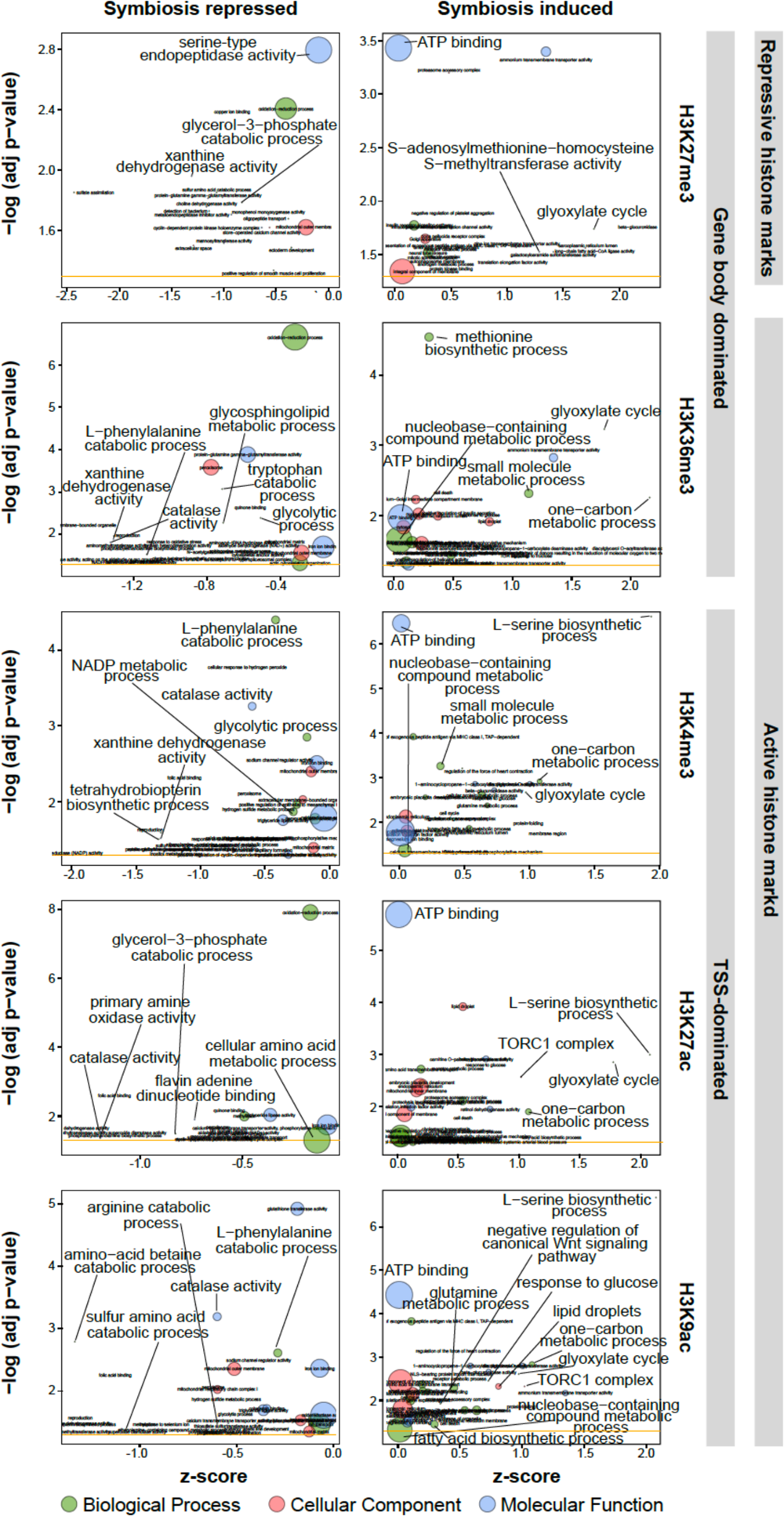
Gene Ontology bubble plots Gene ontology (GO): biological process (BP), cellular component (CC) and molecular function (MF) of all five histone modifications (H3K27me3, H3K36me3, H3K4me3, H3K27ac and H3K9ac) associated genes which are repressed and induced in symbiosis. Y-axis is negative of log adjusted p-value, x-axis is z score and area of the circle is the number of genes in particular category.

### Supplementary File – Extended Methods

Chromatin Immunoprecipitation (ChIP) protocols have been primarily optimized for human, mice and plant cell studies. The work presented here, and the resulting products, are the first attempts at using ChIP on a symbiotic cnidarian in order to understand the histone mechanisms of corals. In a recent publication ^6^, a summarized version of the protocol was published. However, the ChIP protocol is sensitive and therefore requires a number of optimization, validation and quality control steps. In the following, we focus on required pre-protocol validations, the optimization and quality control steps taken pre-IP, leading to the final protocol as applied.

### Validation of experimental concept

#### Histone sequence conservation

While histone modifications are highly conserved across eukaryotes, there has been evidence of histone variants ^73^. Although these variants do not show altered function of the histone in the nucleosome (i.e. packaging of DNA), a potential change in base pair could hinder the binding of commercial anitbodies and, if the target base pair is the modified one, indicate that the modification of interest is not conserved.

The main part of interest is the conservation of the N-terminal tail of histone 3, which carries the modifications investigated in this study. As expected, histone tails are of Aiptasia are highly conserved and align to other plant, animal and fungi sequences (Supp. SF1. Fig. 1). Additionally, we also aligned human histone variants H3.2 and H3.3 to Aiptasia histone models and found continuous conservation of amino acids (98.2% and 100% positives, respectively), indicating that Aiptasia also carries various variant forms of histone 3.

**SF1. Fig. 1.**
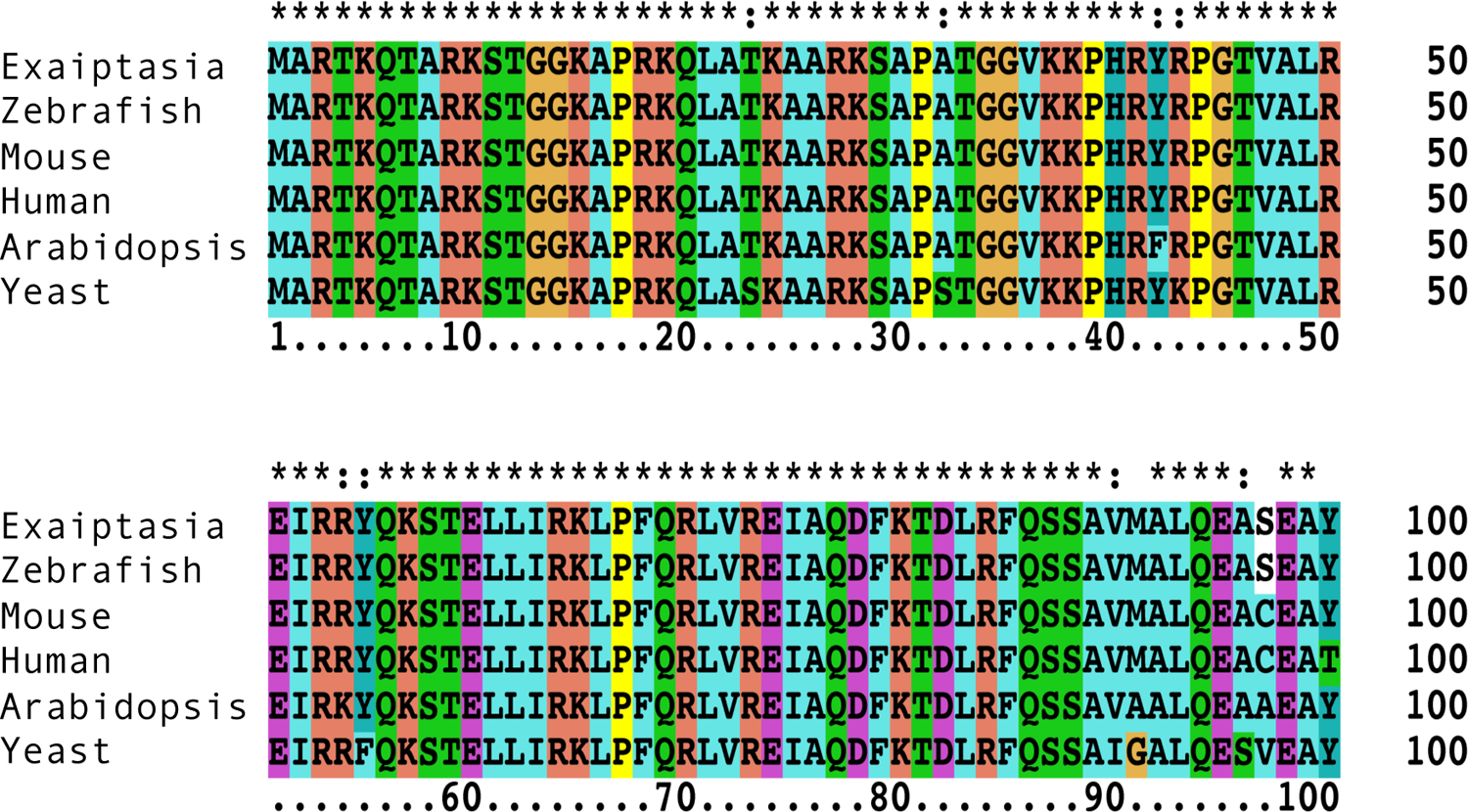
Sequences alignment of histone 3 (H3) across species. Conservation of amino acid bases is consistent across organisms, particularly across the N-terminal tail (adapted from *Li et al., 2018*)

The high conservation of the amino acid sequence, particularly on and around sites of interest such as lysine 4 and 9, indicates that the epitope of commercially available antibodies is present and should be detectable. Interestingly, there appears to be a difference in amino acid between organisms at the 98^th^ position; in Exaiptasia and Zebrafish, the sequence carries serine (S) while mice and humans have cysteine (C). Since these positions are in the fold motif of the histone, no modifications occur there. Hence, alterations in this region are not of concern for the purpose of this study.

#### Antibody validation through Western blot

The success of ChIP and its subsequent sequencing is heavily dependent on the antibody quality. Thus, it is important to validate their affinity and sensitivity in order to be used in ChIP studies. Commerical antibodies for H3K4me3 (ab8580, Abcam), H3K27me3 (ab6002, Abcam), H3K27ac (ab4729, Abcam), H3K36me3 (ab9050, Abcam) and H3K9ac (ab4441, Abcam) were validated for use in Exaiptasia. Total protein extraction and western blot was conducted as described in Li et al (2018). The western blot indicated that all antibodies detected proteins in the expected range, except for H3K4me3, which was highly unspecific (Supp. SF1. Fig. 2).

**SF1. Fig. 2.**
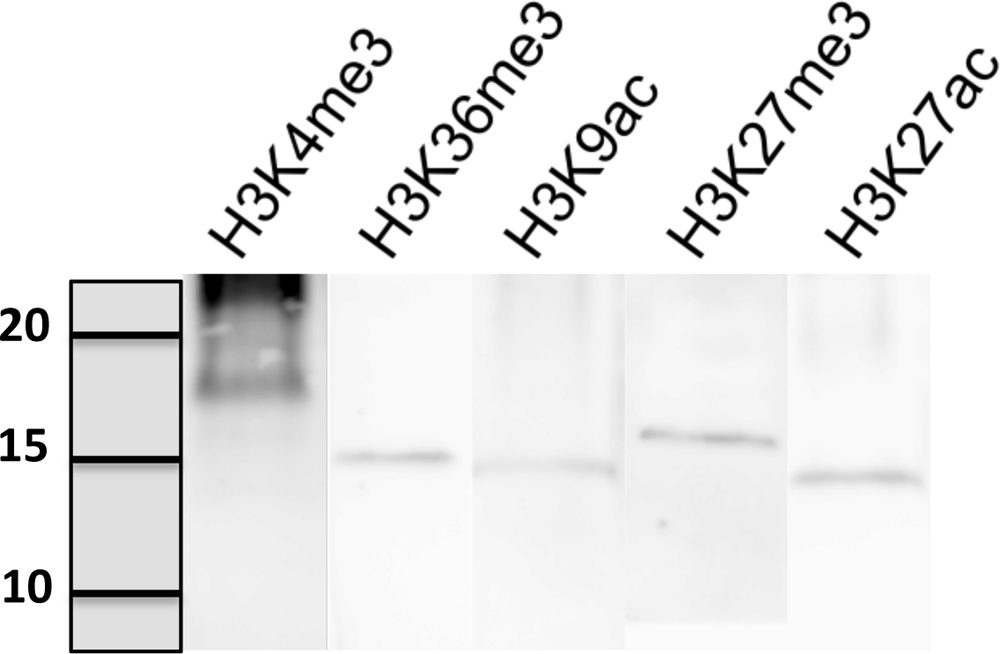
Western blot of histone specific antibodies on total protein content of Aiptasia. Histone proteins lie in a range of 15-20kDa in size.

These results indicated that effective ChIP could only be expected from 4 out 5 antibodies. However, due to the sensitivity of the ChIP protocol, the final validation of the ChIP protocol only occurs one sequenced data is analyzed.

#### ChIP-Seq protocol

Most ChIP protocols are conducted using individually optimized buffers, depending on the type of cells under investigation. Further research into custom buffers versus kits revealed that the customized steps mostly occur primarily prior to the immunoprecipitation (IP). After antibody incubation, wash and clean up steps follow similar principles across protocols; three washes with increasing salinity followed by DNA clean up and elution. In hopes of streamlining future attempts at ChIP-seq in other cnidarians, especially corals, we opted to optimize pre-IP steps to the point that kits could be confidently used thereafter. The optimization steps described here are based on protocols provided by Valerio Orlando Lab (King Abdullah University of Science and Technology, Saudi Arabia) and Moussa Benhamed Lab (Universite Paris-Saclay, France). After lab optimization steps were established the final protocol was executed.

#### Optimization of protocol

##### Pre-IP Buffers

Trial and optimization resulted in two pre-IP buffers being used: the fixation buffer and nucleic preparation buffer. The buffers were adapted from pers. comms. *Valerio Orlando and Schwaiger et al.* (2014), respectively.

#### Fixation buffer

| Chemical | Final concentration |
| --- | --- |
| 1M Hepes-KOH 7.5 pH | 50mM |
| 5M NaCl | 100mM |
| 0.5M EDTA | 1mM |
| 0.5M EGTA | 0.5mM |
| 37% Formaldehyde | 1% |

Add dH<sub>2</sub>O to fill volume

#### Nucleic preparation buffer

| Chemical | Final Concentration |
| --- | --- |
| 1M Hepes-KOH 7.5 pH | 50mM |
| 5M NaCl | 140mM |
| 0.5M EDTA | 1mM |
| 50% Glycerol | 10% |
| 10% Triton 100X | 0.25% |
| 100X PIC | 1X |

Add dH<sub>2</sub>O to fill volume

#### Sonication time

ChIP-seq requires fragment sizes between 100-600 bp. Ideally, shearing fixed histone-DNA should result in around 200-300bp, since one nucleosome packs around 220-250 bp. Because different types of tissue may behave differently during sonication, it’s important to test the efficiency of time series. We determined that, with 1% formaldehyde fixation for 15 minutes, the optimum sonication time was 15 cycles (15 sec ON, minimum 30 sec cooling) to ensure fragmentation to 200-500 bp.

#### ChIP-Seq protocol

We used the Zymo-Spin ChIP Kit (Zymo Research, Irvine, CA) to conduct histone bound chromatin extraction, with minor adjustments to the manufacturer’s protocol. The experiment was conducted on three biological replicates, each consisting of two symbiotic anemones. The following is a detailed explanation of pre-IP steps modified and adjusted for Aiptasia (Supp. SF1. Fig 3):

1. Anemones were spun down and excess water was removed, followed by a quick rinse in 1x PBST (phosphate-buffered saline with 0.1% triton).
2. Anemones were fixed in formaldehyde buffer containing 1% FA for 15 minutes at room temperature
3. Fixation was stopped by adding 1/20 of the volume 2.5M glycine *All following steps until elution of DNA should be conducted on ice*
4. Remove solution and wash anemones in cold 1x PBS
5. Suspend anemones in Nucleic preparation buffer and transfer into a douncer for homogenization. Two anemones were crushed at the same time to produce one biological replicate
6. Transfer homogenized tissue into eppendorf and spin for 5min at 500g to collect cellular debri and larger fragments at the bottom of the tube
7. Take the supernatant and transfer to a clean eppendorf.
8. Spin down and collect nuclei in 4°C at 2000g for 10min
9. Resuspend nuclei in Chromatin shearing buffer provided in the kit.
10. Take a sample of your nuclei and dry on glass slide with DAPI staining. Confirm the presence of intact nuclei under the microscope.
11. Sonicate remaining sample for 15 cycles (15 sec ON, 30 sec cooling).
12. Proceed with IP, wash and elute as described in manufacturer’s protocol.

A corresponding input control was maintained for each of the three biological replicates generated. DNA fragment quality and quantity were confirmed using High Sensitivity DNA Reagents (Agilent Technologies, California, United States) on a bioanalyzer. Upon fragment DNA and fragment size validation, ChIP libraries were constructed using NEBNext ChIP-Seq Library Prep Master Mix Set (NEB, Ipswich, MA).

**SF1. Fig. 3.**
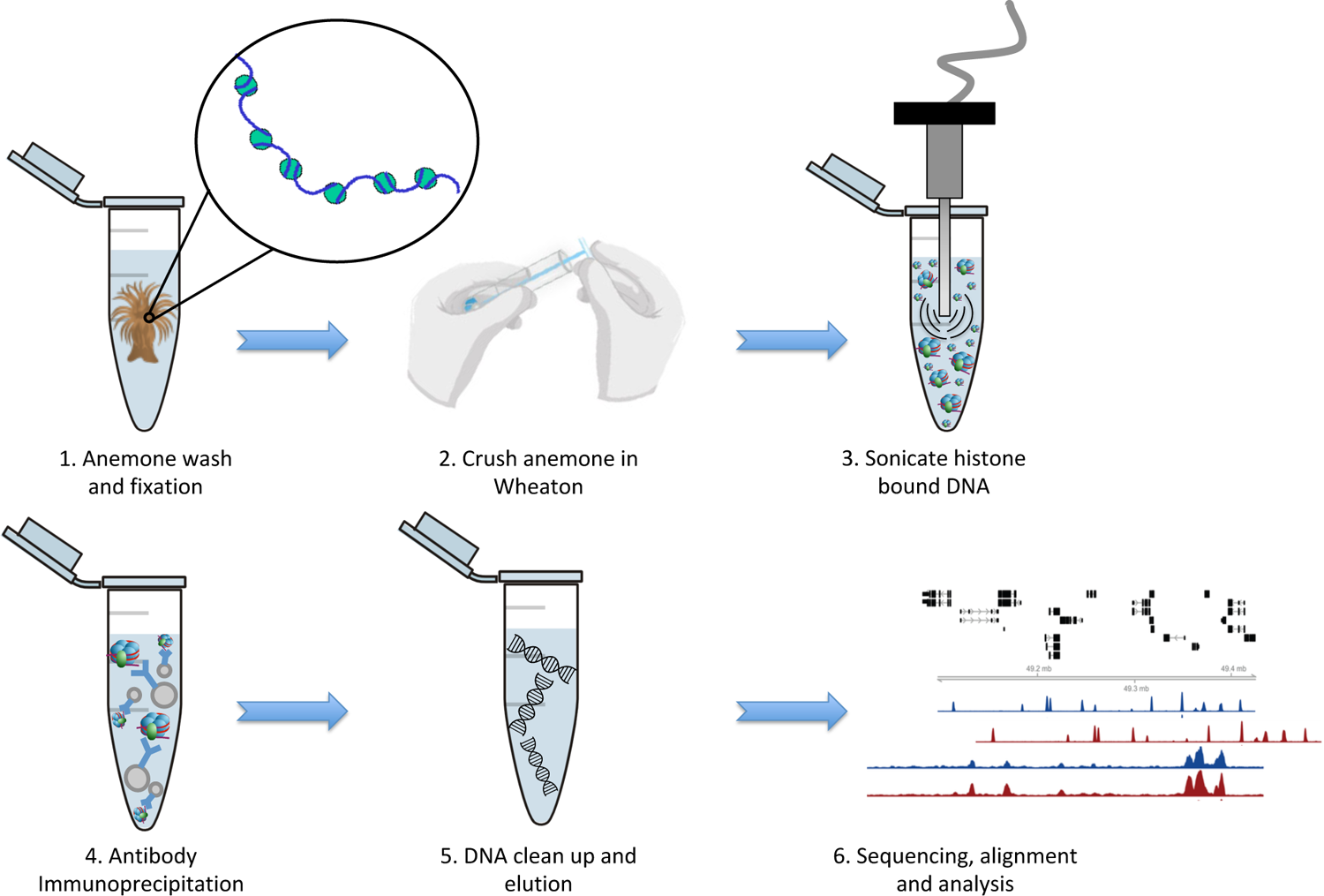
Schematic representation of ChIP-seq protocol using Aiptasia. Detailed description of each step can be found in section III.2.2.2. Only step 1 to 3 were customized for *Aiptasia*; step 4 to 5 were conducted as per manufacturer’s protocol.

## References

1. Dubinsky, Z. & Stambler, N. Coral reefs: an ecosystem in transition. (Springer Netherlands, 2011).

2. Weis, V. M. & Allemand, D. What Determines Coral Health? Science 324, 1153–1155 (2009).

3. Hughes, T. P. et al. Coral reefs in the Anthropocene. Nature 546, 82–90 (2017).

4. Hoegh-Guldberg, O. Climate change, coral bleaching and the future of the world’s coral reefs. Mar. Freshw. Res. 50, 839–866 (1999).

5. Davy, S. K., Allemand, D. & Weis, V. M. Cell Biology of Cnidarian-Dinoflagellate Symbiosis. Microbiol. Mol. Biol. Rev. 76, 229–261 (2012).

6. Li, Y. et al. DNA methylation regulates transcriptional homeostasis of algal endosymbiosis in the coral model Aiptasia. Sci. Adv. 4, eaat2142 (2018).

7. Mazmanian, S., Kasper, D. The love–hate relationship between bacterial polysaccharides and the host immune system. Nat Rev Immunol 6, 849–858 (2006).

8. Cerf-Bensussan, N., Gaboriau-Routhiau, V. The immune system and the gut microbiota: friends or foes?. Nat Rev Immunol 10, 735–744 (2010).

9. Cui, G. et al. Host-dependent nitrogen recycling as a mechanism of symbiont control in Aiptasia. PLOS Genet. 15, e1008189 (2019).

10. Bierne, H. Cross Talk Between Bacteria and the Host Epigenetic Machinery. in Epigenetics and Human Health (eds. W. Doerfler & J. Casadesús) 113–158 (Springer International Publishing, 2017).

11. Kimura, H. Histone modifications for human epigenome analysis. J. Hum. Genet. 58, 439– 445 (2013).

12. Zhou, J. et al. Genome-wide profiling of histone H3 lysine 9 acetylation and dimethylation in Arabidopsis reveals correlation between multiple histone marks and gene expression. Plant Mol. Biol. 72, 585–595 (2010).

13. Pfluger, J. & Wagner, D. Histone modifications and dynamic regulation of genome accessibility in plants. Curr. Opin. Plant Biol. 10, 645–652 (2007).

14. Wolffe, A. P., Wong, J. & Pruss, D. Activators and repressors: making use of chromatin to regulate transcription. Genes Cells 2, 291–302 (1997).

15. Barno, A. R., Villela, H. D. M., Aranda, M., Thomas, T., & Peixoto, R. S. Host under epigenetic control: A novel perspective on the interaction between microorganisms and corals. BioEssays 43, e2100068 (2021).

16. Nagymihály, M. et al. Ploidy-dependent changes in the epigenome of symbiotic cells correlate with specific patterns of gene expression. Proc. Natl. Acad. Sci. U. S. A. 114 (17), 4543– 4548 (2017).

17. Alonso, C., Ramos-Cruz, D. & Becker, C. The role of plant epigenetics in biotic interactions. New Phytol. 221, 731–737 (2019).

18. Negri, I. & Jablonka, E. Editorial: Epigenetics as a deep intimate dialogue between host and symbionts. Front. Genet. 7, 1–3 (2016).

19. Luger, K., Mäder, A. W., Richmond, R. K., Sargent, D. F. & Richmond, T. J. Crystal structure of the nucleosome core particle at 2.8 Å resolution. Nature 389, 251–260 (1997).

20. Olins, D. E. & Olins, A. L. Chromatin history: our view from the bridge. Nat. Rev. Mol. Cell Biol. 4, 809–814 (2003).

21. Jan, B. et al. Nucleosomes, linker DNA, and linker histone form a unique structural motif that directs the higher-order folding and compaction of chromatin. Proc. Natl. Acad. Sci. 95, 14173–14178 (1998).

22. Murray, K. The Occurrence of iε-N-Methyl Lysine in Histones. Biochemistry 3, 10–15 (1964).

23. Paik, W. K. & Kim, S. E-N-dimethyllysine in histones. Biochem. Biophys. Res. Commun. 27, 479–483 (1967).

24. Hempel, K., Lange, H. W. & Birkofer, L. ∈-N-Trimethyllysin, eine neue Aminosäure in Histonen. Naturwissenschaften 55, 37 (1968).

25. Bernstein, B. E. et al. Genomic Maps and Comparative Analysis of Histone Modifications in Human and Mouse. Cell 120, 169–181 (2005).

26. Suzuki, T., Kondo, S., Wakayama, T., Cizdziel, P. E. & Hayashizaki, Y. Genome-Wide Analysis of Abnormal H3K9 Acetylation in Cloned Mice. PLoS One 3, e1905 (2008).

27. Nishida, H. et al. Histone H3 acetylated at lysine 9 in promoter is associated with low nucleosome density in the vicinity of transcription start site in human cell. Chromosom. Res. 14, 203–211 (2006).

28. Roh, T.-Y., Cuddapah, S. & Zhao, K. Active chromatin domains are defined by acetylation islands revealed by genome-wide mapping. Genes Dev. 19, 542–552 (2005).

29. Zhang, Y. & Reinberg, D. Transcription regulation by histone methylation: interplay between different covalent modifications of the core histone tails. Genes Dev. 15, 2343–2360 (2001).

30. Rea, S. et al. Regulation of chromatin structure by site-specific histone H3 methyltransferases. Nature 406, 593–599 (2000).

31. Fuks, F. et al. The Methyl-CpG-binding Protein MeCP2 Links DNA Methylation to Histone Methylation. J. Biol. Chem. 278, 4035–4040 (2003).

32. Lunyak, V. V. et al. Corepressor-Dependent Silencing of Chromosomal Regions Encoding Neuronal Genes. Science 298, 1747–1752 (2002).

33. Cedar, H. & Bergman, Y. Linking DNA methylation and histone modification: Patterns and paradigms. Nat. Rev. Genet. 10, 295–304 (2009).

34. Zhu, J. & Thompson, C. B. Metabolic regulation of cell growth and proliferation. Nat. Rev. Mol. Cell Biol. 20, 436–450 (2019).

35. Dai, Z., Ramesh, V. & Locasale, J. W. The evolving metabolic landscape of chromatin biology and epigenetics. Nat. Rev. Genet. 21, 737–753 (2020).

36. Sebastian, B. et al. The genome of Aiptasia, a sea anemone model for coral symbiosis. Proc. Natl. Acad. Sci. U. S. A. 112 (38), 11893–11898 (2015).

37. Saksouk, N., Simboeck, E. & Déjardin, J. Constitutive heterochromatin formation and transcription in mammals. Epigenetics Chromatin 8, 3 (2015).

38. Suganuma, T. & Workman, J. L. Crosstalk among Histone Modifications. Cell 135, 604– 607 (2008).

39. Valouev, A. et al. Determinants of nucleosome organization in primary human cells. Nature 474, 516–520 (2011).

40. Mani, S. Regulation of Host Chromatin by Bacterial Metabolites. in *Chromatin Signaling and Diseases* 423–442 (Elsevier Inc., 2016).

41. Alenghat, T. et al. Histone deacetylase 3 coordinates commensal-bacteria-dependent intestinal homeostasis. Nature 504, 153–157 (2013).

42. Chang, P. V., Hao, L., Offermanns, S. & Medzhitov, R. The microbial metabolite butyrate regulates intestinal macrophage function via histone deacetylase inhibition. Proc. Natl. Acad. Sci. U. S. A. 111 (6), 2247–2252 (2014).

43. Weizman, E. & Levy, O. The role of chromatin dynamics under global warming response in the symbiotic coral model Aiptasia. *Commun*. Biol. 2, 282 (2019).

44. Schwaiger, M. et al. Evolutionary conservation of the eumetazoan gene regulatory landscape. Genome Res. 24, 639–650 (2014).

45. Young, M. D. et al. ChIP-seq analysis reveals distinct H3K27me3 profiles that correlate with transcriptional activity. Nucleic Acids Res. 39, 7415–7427 (2011).

46. Huang, C. & Zhu, B. Roles of H3K36-specific histone methyltransferases in transcription: antagonizing silencing and safeguarding transcription fidelity. Biophys. Reports 4, 170–177 (2018).

47. Barski, A. et al. High-Resolution Profiling of Histone Methylations in the Human Genome. Cell 129, 823–837 (2007).

48. Choi, J., Lyons, D. B., Kim, M. Y., Moore, J. D. & Zilberman, D. DNA Methylation and Histone H1 Jointly Repress Transposable Elements and Aberrant Intragenic Transcripts. Mol. Cell 77, 310–323.e7 (2020).

49. Sanz, L. A. et al. A mono-allelic bivalent chromatin domain controls tissue-specific imprinting at Grb10. EMBO J. 27, 2523–2532 (2008).

50. Lee, T. I. & Young, R. A. Transcription of eukaryotic protein-coding genes. Annu. Rev. Genet. 34, 77–137 (2000).

51. Kornberg, R. D. The molecular basis of eukaryotic transcription. Proc. Natl. Acad. Sci. U. S. A. 104, 12955–12961 (2007).

52. Kim, T. K. et al. Trajectory of DNA in the RNA polymerase II transcription preinitiation complex. Proc. Natl. Acad. Sci. U. S. A. 94, 12268–12273 (1997).

53. Charlet, J. et al. Bivalent Regions of Cytosine Methylation and H3K27 Acetylation Suggest an Active Role for DNA Methylation at Enhancers. Mol. Cell 62, 422–431 (2016).

54. Lawrence, M., Daujat, S. & Schneider, R. Lateral Thinking: How Histone Modifications Regulate Gene Expression. Trends Genet. 32, 42–56 (2016).

55. Minarovits, J., Banati, F., Szenthe, K. & Niller, H. H. Epigenetic Regulation. Adv. Exp. Med. Biol. 879, 1–25 (2016).

56. Wade, J. T. & Grainger, D. C. Spurious transcription and its impact on cell function. Transcription 9, 182–189 (2018).

57. Neri, F. et al. Intragenic DNA methylation prevents spurious transcription initiation. Nature 543, 72–77 (2017).

58. Miller, J. L. & Grant, P. A. The role of DNA methylation and histone modifications in transcriptional regulation in humans. Subcell. Biochem. 61, 289–317 (2013).

59. Hoffmeister, M. & Martin, W. Interspecific evolution: microbial symbiosis, endosymbiosis and gene transfer. Environ. Microbiol. 5, 641–649 (2003).

60. Douglas, A. E. The microbial dimension in insect nutritional ecology. Funct. Ecol. 23, 38– 47 (2009).

61. Sunagawa, S. et al. Generation and analysis of transcriptomic resources for a model system on the rise: the sea anemone Aiptasia pallida and its dinoflagellate endosymbiont. BMC Genomics 10, 258 (2009).

62. Liew, Y. J. et al. Epigenome-associated phenotypic acclimatization to ocean acidification in a reef-building coral. Sci. Adv. 4, eaar8028 (2018).

63. : Andrews, S. FastQC: a quality control tool for high throughput sequence data. Babraham Bioinforma. (2010).

64. Bolger, A. M., Lohse, M. & Usadel, B. Trimmomatic: a flexible trimmer for Illumina sequence data. Bioinformatics 30, 2114–2120 (2014).

65. Langmead, B., Trapnell, C., Pop, M. & Salzberg, S. L. Ultrafast and memory-efficient alignment of short DNA sequences to the human genome. Genome Biol. 10, R25 (2009).

66. Zhang, Y. et al. Model-based Analysis of ChIP-Seq (MACS). Genome Biol. 9, R137 (2008).

67. Jalili, V., Matteucci, M., Masseroli, M. & Morelli, M. J. Using combined evidence from replicates to evaluate ChIP-seq peaks. Bioinformatics 31, 2761–2769 (2015).

68. Yu, G., Wang, L. G. & He, Q. Y. ChIP seeker: An R/Bioconductor package for ChIP peak annotation, comparison and visualization. Bioinformatics 31, 2382–2383 (2015).

69. Alexa, A., Rahnenführer, J. & Lengauer, T. Improved scoring of functional groups from gene expression data by decorrelating GO graph structure. Bioinformatics 22, 1600–1607 (2006).

70. Freese, N. H., Norris, D. C. & Loraine, A. E. Integrated genome browser: visual analytics platform for genomics. Bioinformatics 32, 2089–2095 (2016).

71. Krueger, F. & Andrews, S. R. Bismark: a flexible aligner and methylation caller for Bisulfite-Seq applications. Bioinformatics 27, 1571–1572 (2011).

72. Ramírez, F., Dündar, F., Diehl, S., Grüning, B. A. & Manke, T. deepTools: a flexible platform for exploring deep-sequencing data. Nucleic Acids Res. 42, W187–W191 (2014).

73. Weber, C. M. & Henikoff, S. Histone variants: dynamic punctuation in transcription. Genes Dev. 28, 672–682 (2014).

74. Roquis, D. et al. The tropical coral Pocillopora acuta displays an unusual chromatin structure and shows histone H3 clipping plasticity upon bleaching. Wellcome Open Res. 9, 195 (2022).

75. Rodriguez-Casariego, J. A. et al. Coral epigenetic responses to nutrient stress: Histone H2A.X phosphorylation dynamics and DNA methylation in the staghorn coral *Acropora cervicornis*. Ecol Evol. 8 (23), 12193–12207 (2018).

